# Phosphatidylglycerol is the lipid donor for synthesis of phospholipid-linked enterobacterial common antigen

**DOI:** 10.1101/2022.10.21.513317

**Authors:** Kinsey N. Morris, Angela M. Mitchell

## Abstract

The gram-negative outer membrane (OM) is an asymmetric bilayer with phospholipids in its inner leaflet and mainly lipopolysaccharide (LPS) in its outer leaflet and is largely impermeable to many antibiotics. In *Enterobacterales* (e.g., *Escherichia, Salmonella, Klebsiella, Yersinia*), the outer leaflet of the OM also contains phosphoglyceride-linked enterobacterial common antigen (ECA_PG_). This molecule consists of the conserved ECA carbohydrate linked to diacylglycerol-phosphate (DAG-P) through a phosphodiester bond. ECA_PG_ contributes to the OM permeability barrier and modeling suggests that it may alter the packing of LPS molecules in the OM. Here, we investigate, in *Escherichia coli* K-12, the reaction synthesizing ECA_PG_ from ECA precursor linked to an isoprenoid carrier to identify the lipid donor that provides the DAG-P moiety to ECA_PG_. Through overexpression of phospholipid biosynthesis genes, we observed alterations expected to increase levels of phosphatidylglycerol (PG) increased synthesis of ECAPG, whereas alterations expected to decrease levels of PG decreased synthesis of ECA_PG_. We discovered depletion of PG levels in strains that could synthesize ECA_PG_, but not other forms of ECA, causes additional growth defects, likely due to the buildup of ECA precursor on the isoprenoid carrier inhibiting peptidoglycan biosynthesis. Our results demonstrate ECAPG can be synthesized in the absence of the other major phospholipids (phosphatidylethanolamine and cardiolipin). Overall, these results conclusively demonstrate PG is the lipid donor for the synthesis of ECA_PG_ and provide a key insight into the reaction producing ECA_PG_. In addition, these results provide an interesting parallel to lipoprotein acylation, which also uses PG as its DAG donor.

**IMPORTANCE:** The outer membrane of gram-negative bacteria is a permeability barrier that prevents the entry of many antibiotics into the cell. However, the pathways responsible for outer membrane biogenesis are potential targets for small molecule development. Here, we investigate the synthesis of a form of enterobacterial common antigen (ECA), ECA_PG_, found in the outer membrane of *Enterobacterales* such as *Escherichia, Salmonella, Klebsiella*, and *Yersinia*. ECA_PG_ consists of the conserved ECA carbohydrate unit linked to diacylglycerol-phosphate—ECA is the headgroup of a phospholipid. The details of the reaction forming this molecule from ECA linked to an isoprenoid carrier are unknown. We determined that the lipid donor that provides the phospholipid moiety to ECA_PG_ is phosphatidylglycerol. Understanding the synthesis of outer membrane constituents such as ECA_PG_ provides the opportunity for the development of molecules to increase outer membrane permeability, expanding the antibiotics available to treat gram-negative infections.

## INTRODUCTION

Gram-negative bacteria have a cell envelope that consists of an inner membrane (IM), aqueous periplasm containing the peptidoglycan cell wall, and an outer membrane (OM). In contrast to the IM and most biological membranes, the OM is an asymmetrical membrane with phospholipids in the inner leaflet and mainly lipopolysaccharide (LPS) in the outer leaflet (1). Beyond its lipid components, the OM is heavily populated with outer membrane proteins and OM lipoproteins (1–3). In addition, the OM contains lower abundance constituents, such as enterobacterial common antigen (ECA) (4), which is found throughout *Enterobacterales*.

The OM presents a strong permeability barrier capable of excluding both large molecules and hydrophobic molecules, including many antibiotics (1, 5, 6). Thus, the biogenesis pathways for OM components are potential targets for the development of small molecules to increase outer membrane permeability and the cell’s susceptibility to antibiotics (7, 8). In fact, several potential antimicrobials targeting biosynthesis of LPS and outer membrane protein biosynthesis have recently been developed (8–15). Although these pathways are found throughout gram-negative bacteria, the biosynthesis of ECA is a pathway that could allow development of small molecules to increase permeability in an order-specific manner, limiting off-target effects on bystander species during treatment.

ECA is an invariant carbohydrate moiety that is found in all *Enterobacterales* (e.g., *Escherichia, Salmonella, Klebsiellla, Shigella, Enterobacter, Yersinia*), with the exception of endosymbionts with greatly reduced genomes (reviewed in (4)). The carbohydrate moiety consists of GlcNAc (*N*-acetylglucosamine), ManNAcA (*N*-acetyl-D-mannosaminuronic acid), and Fuc4NAc (4-acetamido-4,6-dideoxy-D-galactose) which form repeating units of →3)-α-Fuc4NAc-(1→4)-β-ManNAcA-(1→4)-α-GlcNAc-(1→ (**Figure 1A**) (16, 17). ECA can be found in three forms: ECA_CYC_, a cyclic form found soluble in the periplasm, ECA_PG_ (ECA phosphoglyceride), a surface-exposed phospholipid form of ECA (see below), and ECA_LPS_, LPS with ECA attached in place of O-antigen (18–25). We have observed that ECA_CYC_ and ECA_PG_ play roles in maintaining the OM permeability barrier in *Escherichia coli* K-12 (26). Modeling through molecular dynamics has suggested that the presence of ECA_PG_ in the outer leaflet of the OM can lead to changes in packing of LPS with more space allotted per molecule (27, 28). In addition, work in *Salmonella enterica* serovar Typhimurium shows that ECA-deficient strains are non-virulent and establish low level persistent infections (29). Beyond its direct role in pathogenesis, exposure to ECA can lead to the development of antibodies that recognize species throughout *Enterobacterales* (30–32), likely playing an important protective or pathogenic role in *Enterobacterales* infection. These antibodies are most easily generated by exposure to rough *Enterobacterales* strains, which have high levels of ECA_LPS_.

**Figure 1:**
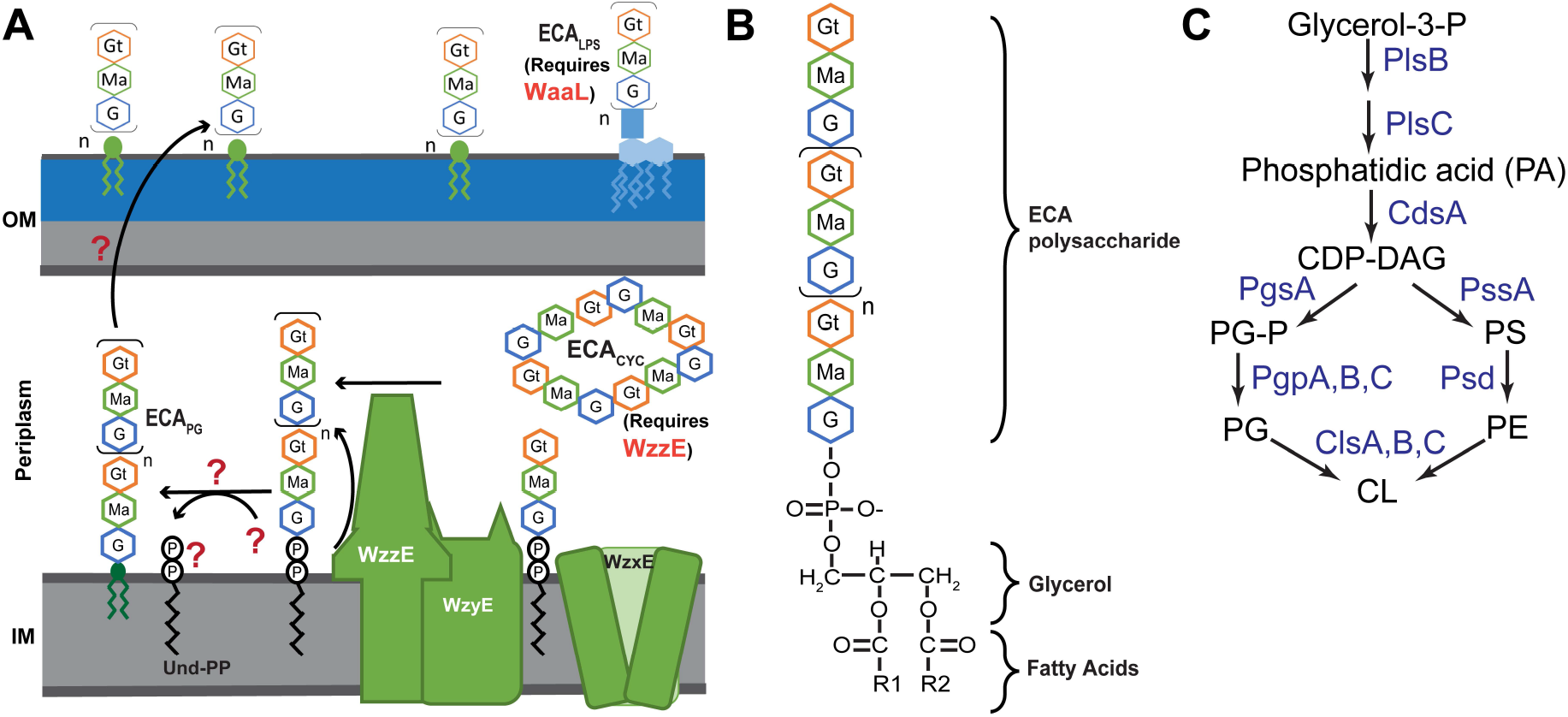
Structure and synthesis of ECA_PG_. **(A)** Subunits of ECA are assembled on the cytoplasmic face of the inner membrane, attached to an isoprenoid carrier, undecaprenyl-pyrophosphate (Und-PP). The subunits are then flipped across the membrane by WzxE and polymerized by WzyE. WzzE controls the chain length of the final ECA molecules. Polymerized ECA can be made into three forms: cyclic ECA (ECA_CYC_), LPS-linked ECA (ECA_LPS_), and ECA_PG_ (see below). ECA_LPS_ synthesis requires the O-antigen ligase, WaaL, for synthesis, while ECA_CYC_ synthesis requires WzzE. The steps and genes required for the formation of ECA_PG_ from polymerized ECA on Und-PP and for its subsequent surface exposure are unknown. **(B)** The structure of the ECA_PG_ glycolipid is shown. The ECA carbohydrate chain is liked to diacylglycerol-phosphate (DAG-P) through a phosphodiester bond. Thus, ECA is linked to the phospholipid backbone in place of a headgroup. **(C)** *E. coli* phospholipid synthesis begins with the addition of acyl chains to glycerol-3-phosphate by PlsB and PlsC forming phosphatidic acid (PA). Then, CdsA activates PA with CDP for form CDP-DAG, the precursor for all phospholipid synthesis. From this point, PgsA and PgpA, PgpB, or PgpC synthesize phosphatidylglycerol (PG), while PssA and Psd synthesize phosphatidylethanolamine (PE). Cardiolipin (CL) is synthesized by ClsA or ClsB from two PG molecules, while ClsC synthesizes CL from one PG and one PE molecule. PS: phosphatidylserine; PG-P: phosphatidylglycerol phosphate.

Given their differences in structure, localization, and phenotypes, it is likely that each form of ECA plays distinct roles in the cells. However, lack of understanding of the steps that differentiate the biosynthesis of the three forms of ECA impedes experimental approaches to investigate their function. ECA is synthesized in a Wzy pathway analogous to that of many O-antigens (**Figure 1A**) (33). ECA trisaccharide units are first assembled on undecaprenyl-pyrophosphate (Und-PP), an isoprenoid carrier, and then flipped to the outer leaflet of the inner membrane by WzxE (4, 34–36). ECA units are polymerized by WzyE with the chain length of the final ECA unit controlled by WzzE (37, 38). Once a polymerized ECA precursor is formed on Und-PP, it can be made into ECA_PG_, ECA_LPS_, or ECA_CYC_. Formation of ECA_CYC_ requires the chain length regulator, WzzE (24), while formation of ECA_LPS_ requires the O-antigen ligase WaaL (18). However, the details of the reaction forming ECA_PG_ and leading to ECA_PG_ surface exposure are completely unknown.

Here, we set out to identify, in *E. coli* K-12, the lipid donor(s) that provides the “phospholipid base” to ECA_PG_. ECA_PG_ consists of the ECA carbohydrate moiety attached to diacylglycerol-phosphate (DAG-P, phosphatidic acid) through a phosphodiester bond (**Figure 1B**) (20, 25). In essence, ECA is a large phospholipid headgroup. It seems biochemically likely that ECA is removed from the isoprenoid carrier freeing Und-PP, as occurs when O-antigen and other forms of ECA are synthesized, and transferred to a specific subset of phospholipids or phospholipid precursors releasing the headgroup in favor of ECA (**Figure 1A**).

In actively growing *E. coli*, the cell’s phospholipid composition is 75% phosphatidylethanolamine (PE), 20% phosphatidylglycerol (PG), and 5% cardiolipin (CL) with the amount of cardiolipin (CL) increasing in stationary phase at the expense of PG (reviewed in (39)). The distribution of phospholipids in the IM is asymmetric with higher PE levels in the inner leaflet than the outer leaflet (40). Phospholipid synthesis begins with the sequential addition of fatty acids to glycerol-3-P by PlsB and PlsC to form phosphatidic acid (PA) (**Figure 1C**) (41). Subsequently, CdsA activates PA through addition of CMP to form CDP-DAG, the precursor for phospholipid biosynthesis (42). From this point, the pathway splits between synthesis of PG and PE. PgsA exchanges CMP for glycerol-3-phosphate to form phosphatidylglycerol phosphate (PG-P) (43). Then, PG-P is dephosphorylated by PgpA, PgpB, or PgpC to form PG (44–46). While PgpA and PgpC are specific to PG-P dephosphorylation (45, 46), PgpB can also dephosphorylate DAG-PP, PA, lyso-PA, and Und-PP (44, 47). For PE synthesis, PssA (Pss) first synthesizes phosphatidylserine (PS) from CDP-DAG (48). Then, Psd decarboxylates PS to form PE (49). *E. coli* has three CL synthase ClsA, ClsB, and ClsC (50–53). ClsA and ClsB form CL from two molecules of PG (50, 54), while ClsC synthesizes CL from one PG molecule and one PE molecule (50). ClsA is mainly responsible for CL synthesis in exponential phase, while all three synthase contribute to CL synthesis in stationary phase.

We examined the effects of alterations in expression of phospholipid biosynthesis on levels of ECA_PG_. Our overexpression data suggest that increasing PG synthesis increases production of ECA_PG_. We determined that the donor for ECA_PG_ synthesis is PG as ECA_PG_ is still synthesized in the absence of CL or PE and depletion of PG leads to inhibited cell growth even in a strain where PG is normally not essential, likely due to the accumulation of ECA precursor on Und-PP inhibiting peptidoglycan biosynthesis. These data are the first to characterize the reaction(s) forming ECA_PG_ and suggest a common approach to use of PG as a lipid donor.

## RESULTS

### Genetic alterations in phospholipid synthesis do not affect ECA_LPS_ levels

The kinetics of biochemical reactions, and so the amount of product produced, often depend on the amount of available substrates for the reaction. For ECA_PG_ to be formed, at least two substrates are necessary: the polymerized ECA precursor on Und-PP and the donor phospholipid or phospholipid precursor. Thus, we hypothesized that changing the availability of the donor lipid would alter the kinetics of the reaction(s) producing ECA_PG_ and so the amount of ECA_PG_ produced. However, changes in phospholipid composition can alter the folding and topology of membrane proteins (reviewed in (55)). As the ECA biosynthesis pathway contains many membrane proteins (4), we asked whether alteration of phospholipid composition would affect ECA synthesis in a manner unrelated to the ECA_PG_ lipid donor. Therefore, we tested the effect of overexpressing genes in phospholipid biosynthesis on levels of ECA_LPS_ to identify any off target effects of phospholipid level alteration on this form of ECA. Using our previously described WGA-staining protocol (56), we found that overexpression of genes in phospholipid biosynthesis caused only minor decreases in the levels of ECA_LPS_ (**Figure 2**). This result is consistent with our previous observation that ECA_LPS_ levels are largely dependent on the availability of WaaL (56) and confirms that ECA can be successfully synthesized with our tested alterations in phospholipid biosynthesis.

**Figure 2:**
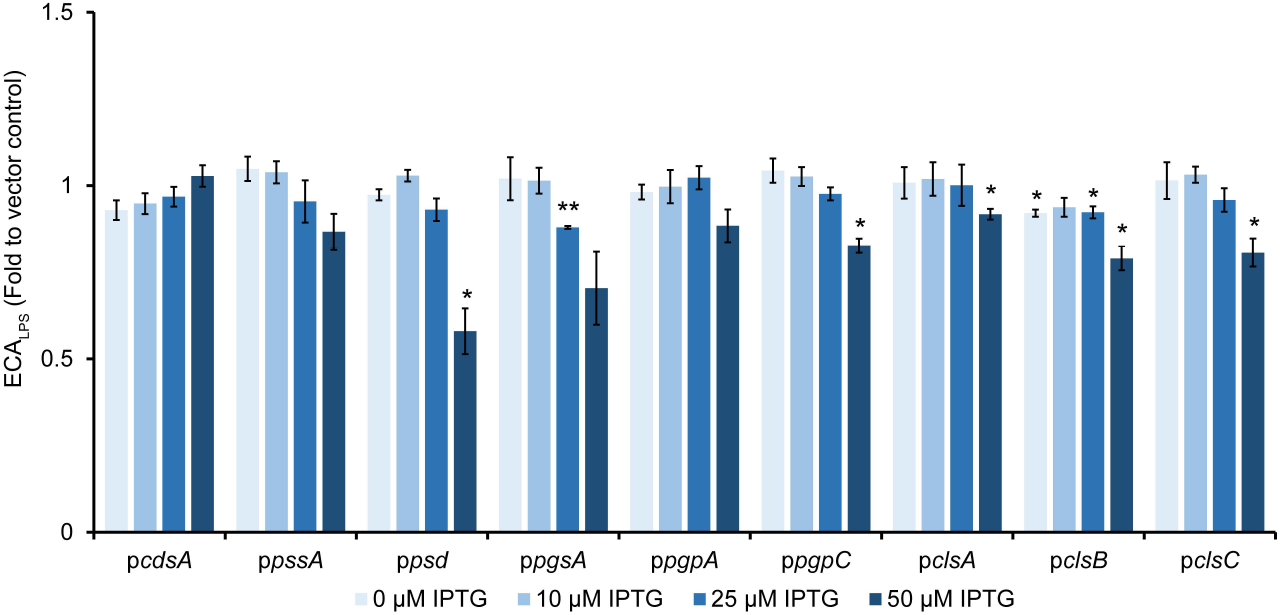
Alterations in phospholipid synthesis cause only minor changes in ECA_LPS_ levels. Levels of ECA_LPS_ were assayed by WGA staining in strains overexpressing the indicated genes in phospholipid biosynthesis. Data are displayed as fold values to the vector control. Only minor decreases and no significant increases in ECA_LPS_ levels were observed. Data are the mean of three biological replicates ± the SEM. * p<0.05 by paired T-test; ** p<0.005 by paired T-test

### The ECA_PG_ lipid donor is not PA

We investigated the effect of phospholipid gene overexpression on the accumulation of ECA_PG_ and on activity of the P*_wec_* promoter region responsible for expression of the *wec* operon that contains the genes necessary for synthesis of the polymerized ECA precursor. We reasoned overexpression of genes before the lipid donor in the phospholipid biosynthesis pathway would increase the amount of the lipid donor and so the amount of ECA_PG_. Conversely, overexpression of genes after the donor or on a different branch of the pathway would decrease the amount of the donor and so the amount of ECA_PG_. Changes in P*_wec_* activity suggest changes to ECA precursor levels that may be occurring, in addition to any changes in lipid donor levels.

The lipid to which ECA is attached in ECA_PG_ is DAG-P and so is most similar to PA. Therefore, we assayed the effect of *cdsA*, which converts PA into CDP-DAG, overexpression under an IPTG inducible promoter on ECA_PG_ levels. To allow specific detection of ECA_PG_, we tested ECA_PG_ levels in a Δ*wzzE* Δ*waaL* strain that only produces ECA_PG_ and not the other two forms of ECA. At all tested IPTG concentrations, we observed a large increase in the level of ECA_PG_ (**Figure 3A**, lanes 2 vs. 3-6). In contrast, there were no significant increases in P*_wec_* activity when *cdsA* was overexpressed (**Figure 3B**). These data indicate that the lipid donor for ECA_PG_ synthesis is CDP-DAG and/or a phospholipid but is not PA or an earlier biosynthetic intermediate.

**Figure 3:**
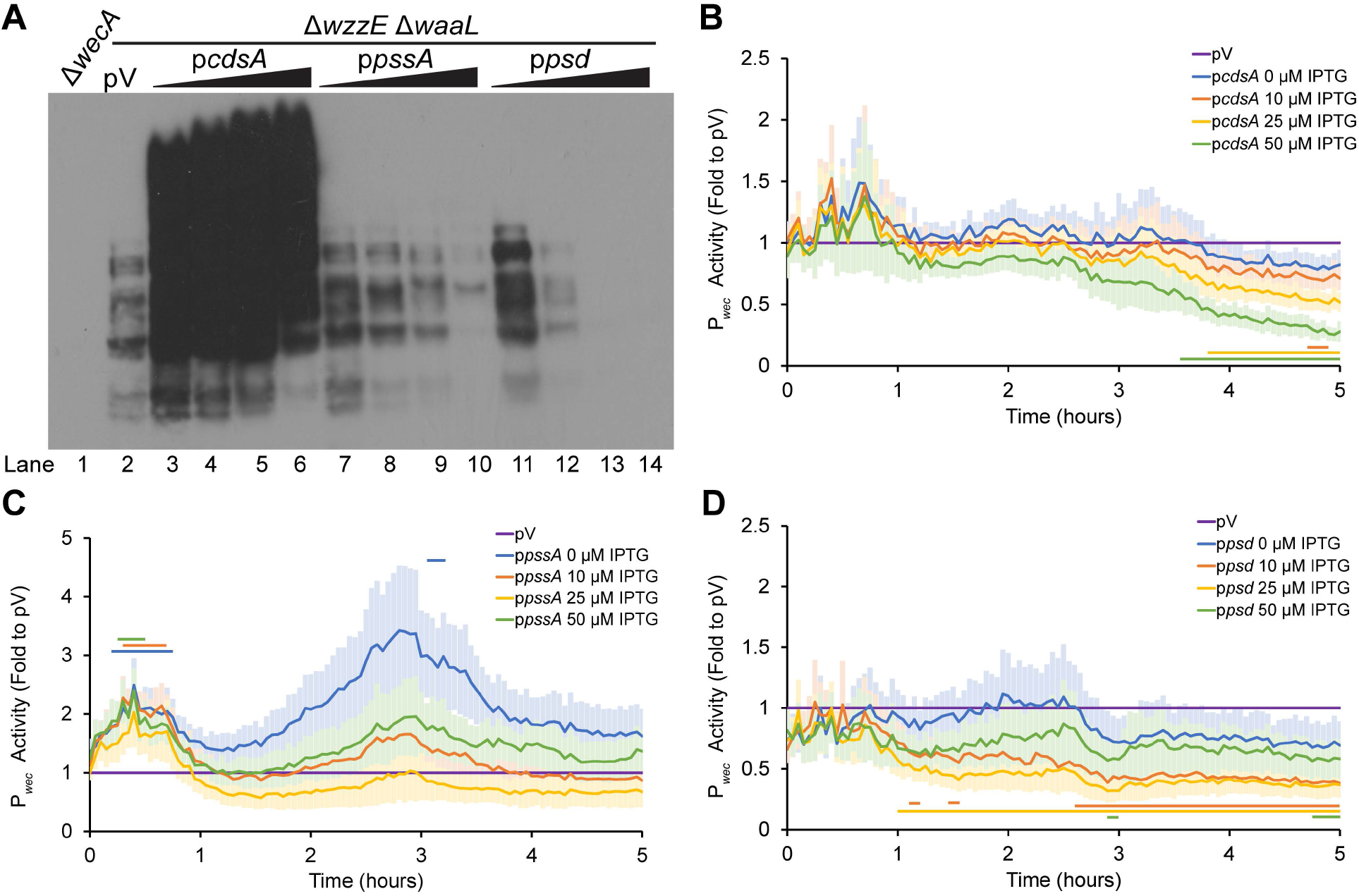
A phospholipid is the substrate for ECA_PG_ synthesis. **(A)** Levels of ECA_PG_ were assayed by immunoblot in a strain that produces only ECA_PG_ and not the other two forms of ECA (Δ*wzzE* Δ*waaL*). The vector control (pV) treated with 50 μM IPTG is compared to strains overexpressing the indicated genes in phospholipid biosynthesis with increasing concentrations of IPTG (0 μM, 10 μM, 25 μM, and 50 μM). A strain with a deletion in *wecA*, the first gene in the ECA biosynthesis pathway serves as a negative control. Overexpression of *cdsA* causes large increases in the amounts of ECA_PG_, while overexpression of *pssA* and *psd* decrease ECA_PG_ levels. Image is representative of three independent experiments. **(B-D)** Strains overexpressing genes in phospholipid biosynthesis and carrying a luciferase reporter for *wec* operon promoter (P*_wec_*) activity were assayed for luminescence and OD600. Data are shown as the fold value of the relative luminescence to the vector control and are the mean from six biological replicates ± the SEM. The empty vector (pV) sample contained the empty vector for phospholipid gene overexpression and the P*_wec_* reporter and was treated with 50 μM IPTG. Horizontal bars: p<0.05 by T-test consistently for three or more time points. **(B)** Strains overexpressing *cdsA* have very similar P*_wec_* activity to that of the empty vector control, with a decrease later in growth. **(C)** Strains overexpressing *pssA* have some increase in P*_wec_* activity. **(D)** Overexpression of *psd* decreases P*_wec_* activity.

### PE synthesis is off the pathway of the lipid donor

To determine whether PE could serve as the donor for ECA_PG_ synthesis, we then overexpressed the genes in PE synthesis pathway, *pssA* and *psd*. Overexpression of these genes caused a decrease in the accumulation of ECA_PG_ in a manner that correlates with the concentration of IPTG (**Figure 3A**, lane 2 vs. 7-10 and 11-14). Overexpression of *pssA* caused some significant increases in P*_wec_* activity at early time points after induction but no decreases in activity (**Figure 3C**). However, *psd* overexpression caused decreased P*_wec_* activity at higher IPTG concentrations (**Figure 3D**), confounding interpretations of decreased ECA_PG_ accumulation observed with *psd* overexpression. Nevertheless, the decreased ECA_PG_ levels without changes in P*_wec_* activity with *pssA* overexpression demonstrate the pathway for synthesis of PE is not involved in synthesis of the ECA_PG_ lipid donor.

### Overexpression suggests the ECA_PG_ lipid donor is part of the PG/CL synthesis pathway

As PE synthesis appeared off pathway for the synthesis of the ECA_PG_ lipid donor, we then examined the effects of gene overexpression in the PG synthesis pathway. As PG is necessary for synthesis of CL, overexpression of these genes would also be expected to increase the synthesis of CL (50, 54). Overexpression of *pgsA*, the first dedicated gene in PG synthesis, caused an increase in ECA_PG_ levels at the highest IPTG concentration we tested (**Figure 4A**, lane 2 vs. 3-6). However, *pgsA* overexpression caused a consistent increase in P*_wec_* activity (**Figure 4B**), making these data difficult to interpret. Overexpression of *pgpA* led to higher levels of ECA_PG_ at all IPTG concentrations assayed (**Figure 4A**, lane 2 vs. 7-10), despite very little significant change in P*_wec_* activity (**Figure 4C**). This result suggests that the lipid donor for ECA_PG_ synthesis may be downstream of PgpA in the biosynthesis pathway.

**Figure 4:**
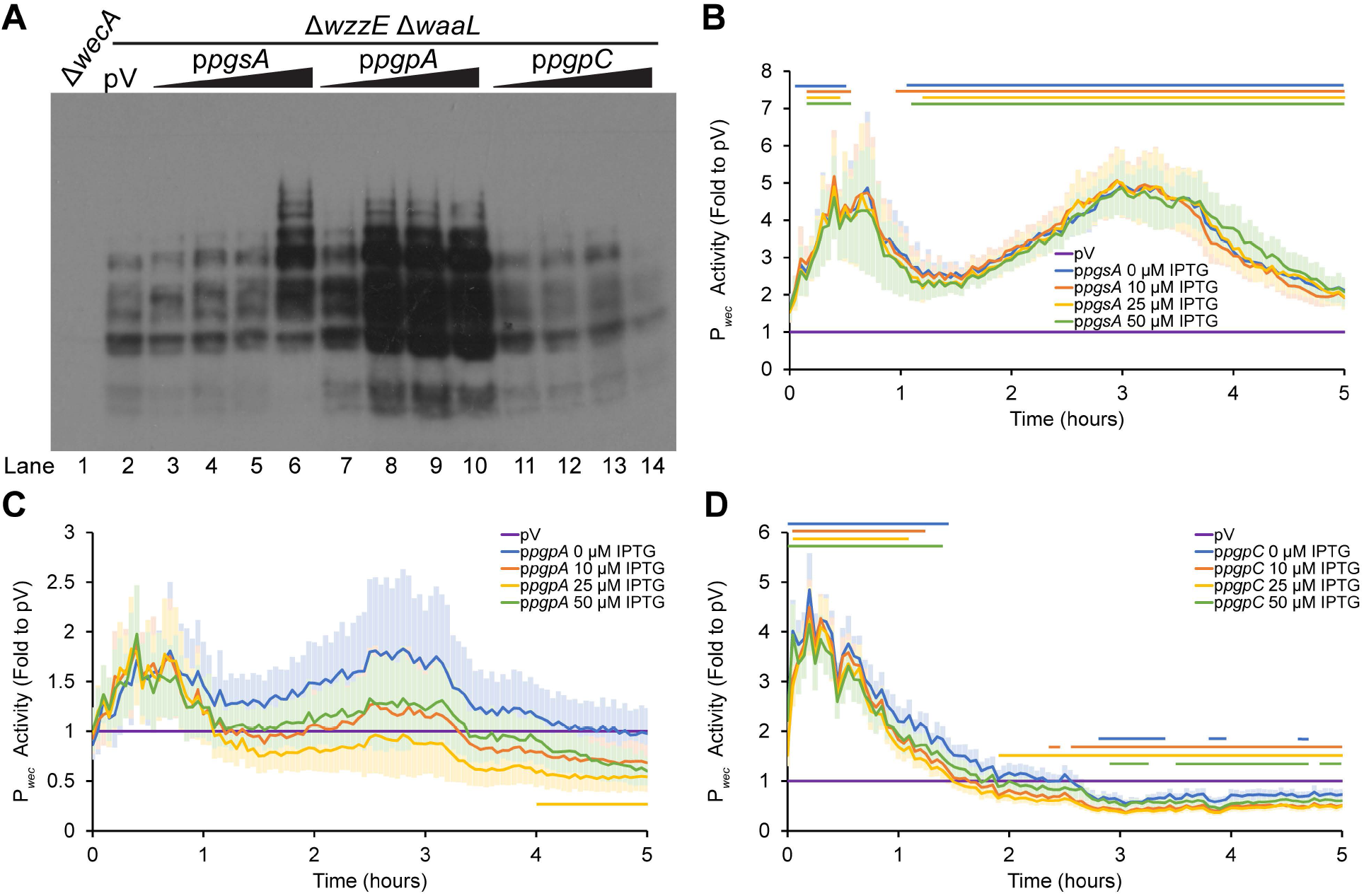
The substrate for ECA_PG_ biosynthesis is in the PG/CL biosynthesis pathway. **(A)** ECA_PG_ levels were assayed by immunoblot analysis in strains overexpressing the indicated genes in phospholipid biosynthesis. The Δ*wzzE* Δ*waaL* strain background allows the production of ECA_PG_ without the other forms of ECA. For phospholipid synthesis gene overexpression, cultures were treated with 0 μM, 10 μM, 25 μM, and 50 μM IPTG. The vector control (pV) was treated with 50 μM IPTG. Δ*wecA* serves as a negative control. Image is representative of three independent experiments. Overexpression of both *pgsA* and *pgpA* increase accumulation of ECA_PG_. Overexpression of *pgpC* causes a small decrease in ECA_PG_ levels. **(B-D)** P*_wec_* activity was assayed in strains overexpressing genes in phospholipid biosynthesis as in Figure 3. The mean ± the SEM for the relative luminescence as a fold value to the vector control is shown for six biological replicates. pV: Empty vector sample carrying the P*_wec_* reporter and the empty vector for the phospholipid synthesis gene overexpression and treated with 50 μM IPTG. Horizontal bars: p<0.05 by T-test consistently for three or more time points. **(B)** Overexpressing *pgsA* consistently increased P*_wec_* activity. **(C)** Overexpression of *pgpA* did not change P*_wec_* activity at most time points and IPTG concentrations. **(D)** Overexpressing *pgpC* causes early increases and later decreases in P*_wec_* activity.

Interestingly, when we overexpressed *pgpC*, we observed the opposite phenotype, a decrease in ECA_PG_ levels at the highest IPTG concentration (**Figure 4A**, lane 2 vs. 11-14). There are two likely explanations for the contradictory results between *pgpA* and *pgpC* overexpression. The first is that the decrease in ECA_PG_ levels with *pgpC* is due to the lower P*_wec_* activity observed several hours after induction of *pgpC* overexpression (**Figure 4D**). The second is due to the relative activities of PgpA, PgpB, and PgpC. Of the three enzymes, PgpA has the highest activity and the most effect on the levels of PG-P and PG (46). Thus, it is possible that overexpression of *pgpC* actually decreases flux through the PG/CL synthesis pathway. We did not overexpress *pgpB*, as PgpB is active on several different substrates including Und-PP (47). Overall, these results suggest that PG or CL may be the donor for ECA_PG_ biosynthesis.

### CL synthesis is downstream from the ECA_PG_ lipid donor

There are three CL synthases, ClsA, ClsB, and ClsC (50–53). When we overexpressed the gene for the CL synthase active in exponentially growing cells, *clsA*, we observed very little change in either ECA_PG_ levels (**Figure 5A**, lane 2 vs. 3-6) or in P*_wec_* activity (**Figure 5B**). This may be due to feedback inhibition of CL on ClsA activity (57). However, when we overexpressed either *clsB* or *clsC*, which are normally active in stationary phase when CL levels increase, we observed a decrease in ECA_PG_ levels that was more severe at higher IPTG concentrations (**Figure 5A**, lane 2 vs. 7-10 and 11-14). Overexpression of *clsB* did not cause significant changes in P*_wec_* activity (**Figure 5C**), while overexpression of *clsC* caused some decreases in P*_wec_* activity at higher IPTG concentrations (**Figure 5D**). These data suggest that CL is downstream of the lipid donor for ECA_PG_ synthesis.

**Figure 5:**
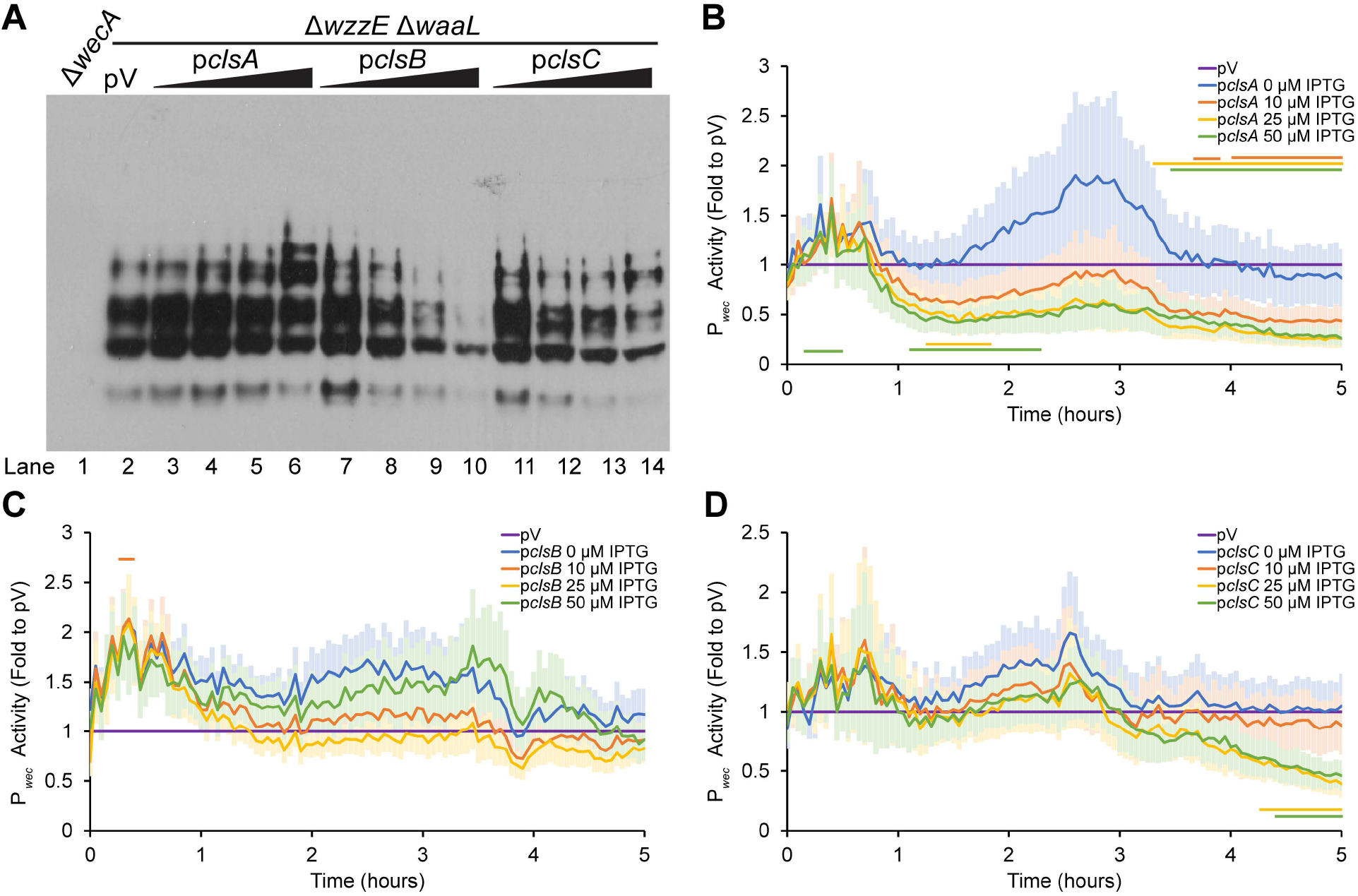
CL synthase overexpression decreases ECA_PG_ levels. **(A)** ECA_PG_ levels were assayed by immunoblot in strains overexpressing genes in CL biosynthesis. The Δ*wzzE*Δ*waaL* strain produces only ECA_PG_ and not the other two forms of ECA. The vector control (pV) was treated with 50 μM IPTG, while overexpression strains were treated with 0 μM, 10 μM, 25 μM, and 50 μM IPTG. The Δ*wecA* strain serves as a negative control. Image is representative of three independent experiments. Overexpression of both *clsB* and *clsC* decreases ECA_PG_ levels while levels of ECA_PG_ are constant with *clsA* overexpression. **(B-D)** The activity of P*_wec_* was assayed as in Figure 3. Data are shown as the fold of the relative luminescence to the vector control (pV) and are the mean of 5 to 6 biological replicates ± the SEM. pV control carried the P*_wec_* reporter and an empty vector for overexpression and was treated with 50 μM IPTG. Horizontal bars: p<0.05 by T-test consistently for three or more time points. **(B)** Overexpression of *clsA* causes some significant increases and decreases in P*_wec_* activity at higher IPTG concentrations. **(C)** Overexpression of *clsB* does not change P*_wec_* activity. **(D)** Overexpression of *clsC* causes some decrease in P*_wec_* activity at higher IPTG concentrations and later time points.

### PG is the substrate for ECA_PG_ biosynthesis

Taken as a whole, our overexpression data point to PG as the lipid donor for ECA_PG_ synthesis: Overexpression of genes expected to increase levels of PE and CL at the expense of PG decreases levels of ECA_PG_, while overexpression of genes expected to increase levels of PG increases levels of ECA_PG_. Therefore, we decided to examine the growth of strains where genes in the PG/CL biosynthesis pathway could be depleted. ECA and peptidoglycan are both synthesized on Und-P (58, 59) and so interruption of ECA biosynthesis can lead to the buildup of Und-PP-linked intermediates, disrupting peptidoglycan biosynthesis (24, 56, 60–64). The severity of the peptidoglycan defect depends on the stage in biosynthesis interrupted. Defects in ECA biosynthesis leading to the accumulation of Und-PP-GlcNAc-ManNAcA (Lipid II^ECA^) produce cell shape defects, envelope permeability, and envelope stress response activation (60, 61, 63). Disruption of the flippases capable of flipping Und-PP-GlcNAc-ManNAcA-Fuc4NAc (Lipid III^ECA^) ECA across the IM or of the ECA polymerase, *wzyE*, is lethal in *E. coli* K-12 (24). We have speculated that, in a strain that makes only ECA_PG_, disruption of the next step in ECA_PG_ biosynthesis would be lethal as well (56). In fact, we have shown that dysregulation of ECA_PG_ biosynthesis in a strain making only ECA_PG_ is lethal (56). Therefore, we hypothesized that loss of the lipid donor for ECA_PG_ biosynthesis would cause peptidoglycan synthesis defects, and so growth phenotypes, specifically when ECA_PG_ is made without the other forms of ECA.

Our depletion strategy involved cloning phospholipid biosynthesis genes under the control of the P_BAD_ promoter, which is arabinose inducible and subject to catabolite repression (65). We then deleted the chromosomal copies of the affected genes while maintaining expression from the plasmid-borne copy in either the wild-type background or the Δ*wzzE* Δ*waaL* strain that produces ECA_PG_ without the other two forms of ECA. Treating cells with arabinose or glucose in the presence of the empty plasmid produced similar growth curves in the wild-type and Δ*wzzE* Δ*waaL* strains (**Figure 6A**). We next examined the effect of depletion of PgpC in a background with the chromosomal copies of *pgpA, pgpB*, and *pgpC* deleted. With depletion, this strain would be expected to have little or no PG or CL and to build up PG-P (46). This depletion was lethal in both the wild-type and Δ*wzzE* Δ*waaL* strains (**Figure 6B**). However, the Δ*wzzE* Δ*waaL* strain ceased growth and began to lyse earlier than the strain with a wild-type background. This suggested that there might be an additional defect in the Δ*wzzE* Δ*waaL* strain beyond that caused by buildup of PG-P.

**Figure 6:**
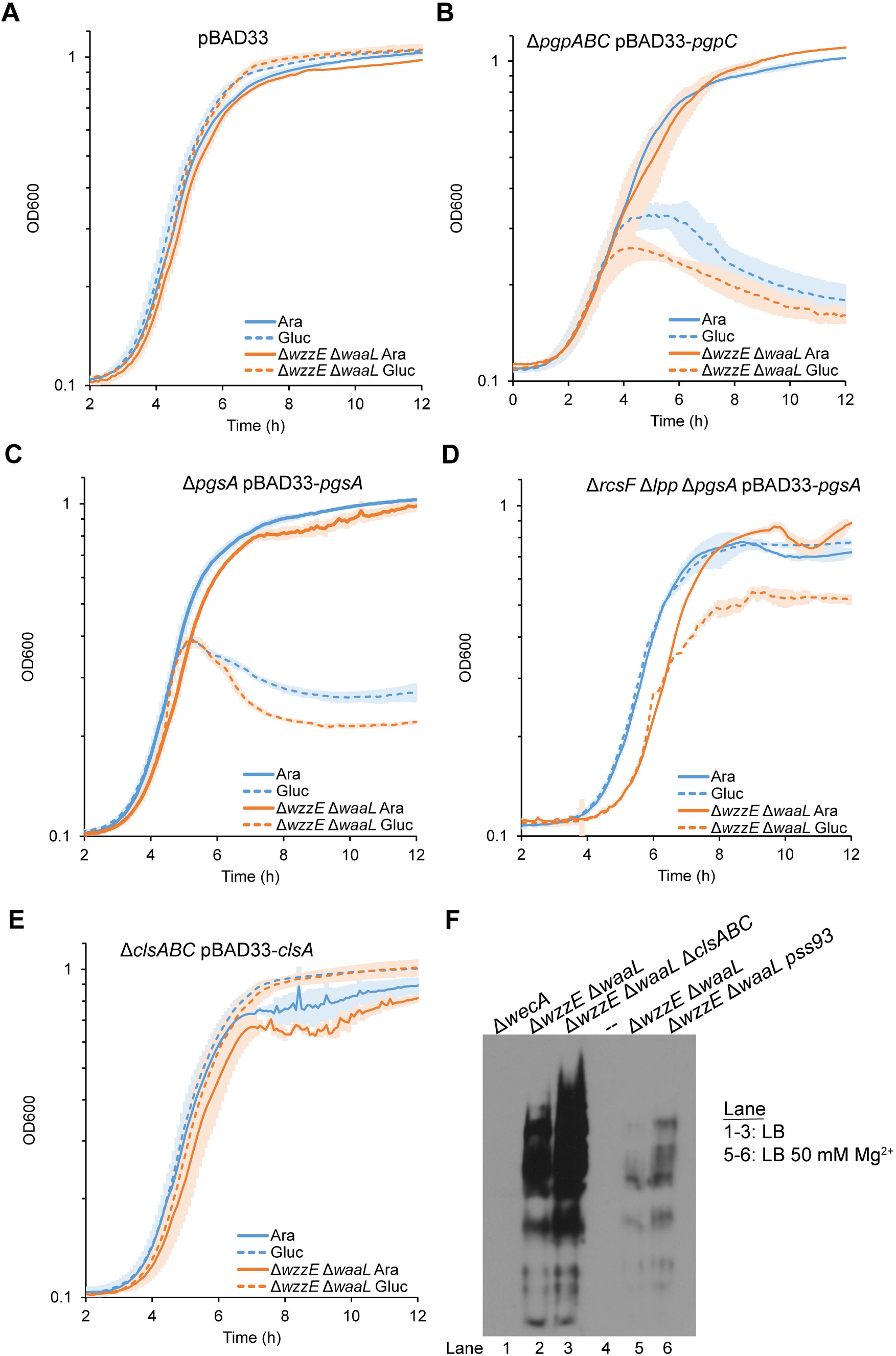
Phosphatidylglycerol is the substrate for the synthesis of ECA_PG_. **(A-E)** Growth curves were performed strains with the indicated genotype alone (blue) or with Δ*wzzE* Δ*waaL* (orange) carrying a plasmid with the P*_BAD_* promoter to investigate disruptions to peptidoglycan biosynthesis caused by accumulation of ECA precursor. Cultures were treated with arabinose to induce expression from the P*_BAD_* promoter (solid lines) or glucose to repress expression (hashed lines). Growth curves were started with a 1:10,000 dilution of cultures grown with arabinose unless otherwise noted. Data are the mean of three biological replicates ± the SEM. **(A)** Strains carrying an empty plasmid showed very similar growth when treated with arabinose or glucose. **(B)** Cultures grown in arabinose were diluted 1:500 for growth curves. Depletion of PgpC in strains with deletions of *pgpA, pgpB*, and *pgpC*caused lysis. However, strains that can make only ECA_PG_ from ECA precursor (Δ*wzzE* Δ*waaL*) lysed earlier in growth than strains that can make all three forms of ECA. **(C)** Depletion of PgsA causes more lysis in the Δ*wzzE*Δ*waaL* strain background than in the wild type background. **(D)** PG is not essential in strains where pressure on lipoprotein maturation are relieved (Δ*rcsF* Δ*lpp*). Depletion of PgsA in this background allowed full growth. However, when this background is combined with Δ*wzzE*Δ*waaL*, cultures in which PgsA depleted cannot finish their growth. **(E)** CL is not essential. Depletion of ClsA in strains with deletions in *cIsA, cIsB*, and *cIsC* did not cause lysis either alone or when combined with Δ*wzzE* Δ*waaL*. **(F)** Immunoblot analysis was used to assay ECA_PG_ levels in strains without CL (Δ*clsABC*) or without PE (*pss93*) Strains lacking PE require high levels of Mg^2+^ for survival, so cultures for lanes 5 and 6 were grown with 50 mM Mg^2+^. Δ*wecA* serves as a negative control. Both strains lacking CL and strains lacking PE retained production of ECA_PG_. Images are representative of two independent experiments.

We then assayed the effect of PgsA depletion as this strain would not build up PG-P and excess CDP-DAG could be funneled into PE. This strain grew longer after depletion and showed less lysis in a wild-type background (**Figure 6C**). The Δ*wzzE* Δ*waaL* strain grew to the same OD as the wildtype background strain but demonstrated quicker and more severe lysis. These results suggested a peptidoglycan defect due to accumulation of ECA precursor on Und-PP.

*pgsA* is essential due to the necessity for PG for lipoprotein processing (66–69). The essentiality of *pgsA* can be circumvented by deletion of *lpp* (66). Lpp (Braun’s lipoprotein) is highly abundant and IM localization is lethal due to crosslinking of the peptidoglycan to the IM (70). Δ*lpp* Δ*pgsA* strains are still temperature sensitive, however (66). This temperature sensitivity can be relieved by inactivating the Rcs stress response (67), which is over-activated by localization of RcsF in the inner membrane (71). Therefore, we built PgsA depletion strains in Δ*rcsF* Δ*lpp* and Δ*rcsF* Δ*lpp* Δ*wzzE* Δ*waaL* backgrounds to test the effect of PgsA depletion in conditions were PG is normally not essential. In the Δ*rcsF* Δ*lpp* background, the PgsA depletion strain grew equally with arabinose or glucose, confirming PG is not essential in this strain (**Figure 6D**). However, the Δ*rcsF* Δ*lpp* Δ*wzzE* Δ*waaL* background strain ceased growth when depleted. These data confirm a second growth defect in this strain beyond that observed in the wild-type background.

Depletion of either PgpABC or PgsA will reduce the level of both PG and CL. Thus, to confirm that the results we observed with depletion were the result of loss of PG and not loss of CL, we assayed the growth of a ClsA depletion strain in a Δ*clsA* Δ*clsB* Δ*clsC* background. CL is not essential and this strain grew fully when depleted in both the wild type and Δ*wzzE* Δ*waaL* strain (**Figure 6E**). These data confirm that it is loss of PG not CL that causes a growth defect with PgsA and PgpABC depletion.

To finalize our conclusion that PG was the lipid donor for the synthesis of ECA_PG_, we attempted to build deletion strains for the PG, PE, and CL synthesis genes in the Δ*wzzE* Δ*waaL* background. Although PE makes up 75% of the membrane in wild-type cells (39), strains with disruption of *pssA* are viable as long as they are maintained with 20 to 50 mM Ca^2+^ or Mg^2+^ (72). We were able to build strains with a *pssA* disruption allele and with Δ*clsA* Δ*clsB* Δ*clsC*. When we assayed these strains, we found that both strains had increased ECA_PG_ levels compared to the Δ*wzzE* Δ*waaL* strain (**Figure 6F**). We also observed a large decrease in ECA_PG_ levels with Mg^2+^ treatment. Consistent with the growth defects we observed in depletion strains (**Figure 6D**), we were not successful in building a Δ*rcsF* Δ*lpp* Δ*wzzE* Δ*waaL* Δ*pgsA* and reviving it from frozen stocks. Together, our data demonstrate that PG is the lipid donor for the synthesis of ECA_PG_.

## DISCUSSION

In this work, we investigated the identity of the lipid donor for synthesis of ECA_PG_. Our data demonstrate through several lines of evidence that this donor is PG and that other phospholipids cannot efficiently substitute when PG is lost. (i) Changes in flux through phospholipid biosynthesis expected to increase PG production increase ECA_PG_ levels, while changes expected to decrease PG production decrease ECA_PG_ levels. (ii) Depletion of PG levels when ECA precursors cannot be made into other forms of ECA inhibits growth, even where PG is otherwise non-essential. (iii) ECA_PG_ is produced in the absence of PE or CL. These results also demonstrate preventing transfer of ECA from Und-PP to the lipid donor causes sufficient peptidoglycan stress to inhibit growth despite the Und-PP released when ECA subunits are polymerized.

In our previous investigations (56), we used this peptidoglycan stress phenotype to identify a system that regulates the production of ECA_PG_. ElyC and ECA_CYC_ work together to post-transcriptionally inhibit the production of ECA during normal growth conditions (56). We envision this system as a feedback pathway that regulates ECA_PG_ production based on levels of ECA_CYC_. We can now add the identity of the lipid donor for ECA_PG_ synthesis to our model of ECA_PG_ synthesis. In our model, polymerized ECA is removed from Und-PP and transferred to the DAG-P portion of a PG molecule, releasing glycerol into the periplasm and Und-PP into the outer leaflet of the IM. The protein(s) responsible for this reaction would be regulated by ElyC in a manner controlled by ECA_CYC_. This model provides a framework for further investigations of the synthesis of ECA_PG_ including: identification of the gene(s) involved in synthesizing ECA_PG_ from PG and the ECA precursor, determining whether the synthesizing enzyme(s) have specific preferences for PG fatty acid composition, the details of the reaction forming ECA_PG_ and of the mechanism of regulation by ElyC and ECA_CYC_, and the pathway leading to ECA_PG_ surface exposure.

In addition to regulation of ECA_PG_ production by ElyC and ECA_CYC_ (56), our results have revealed other mechanisms through which ECA_PG_ production can be regulated, specifically in strains lacking ECA_CYC_ and so full activity of the ElyC-ECA_CYC_ pathway. We relied in our overexpression experiments on the ability of ECA_PG_ production to be altered by lipid donor availability and found the changes in ECA_PG_ production to be robust, with both quite large increases and decreases in ECA_PG_ levels (for example see **Figure 3A**, lane 6 vs. 10). These data are especially interesting given that the ratio of PG to CL varies based on growth phase with the amount of CL increasing in stationary phase (39, 73). Our data suggest that this change may have a direct effect on the synthesis of ECA_PG_.

Beyond regulation by substrate availability, we observed several changes in phospholipid synthesis gene expression that led to changes in the activity of the promoter region of the *wec* operon that encodes the genes necessary to synthesize the ECA precursor. The largest of these changes was the 4-to 5-fold increase in P*_wec_* activity we observed when we overexpressed *pgsA* (**Figure 4B**). These data demonstrate an interplay between phospholipid synthesis and ECA synthesis and suggest that there may be feedback from membrane conditions that affects ECA synthesis. We also observed a large decrease in ECA_PG_ levels when cells were grown with a high concentration of Mg^2+^ (**Figure 6F**). This decrease was present in both the strain with normal phospholipid composition and the strain lacking PE. It is possible that this decrease could be due to changes in Mg^2+^ sensing transcriptional regulation pathways, due to changes in OM order due to increase bridging between LPS molecules, or to changes in the activity of enzymes in the ECA synthesis pathway. Experiments investigating these phenotypes are ongoing in our lab.

It is interesting to compare the synthesis of ECA_PG_ to another glycolipid with a very similar structure, membrane protein intergrase (MPIase) (reviewed in (74)). MPIase consists of the same carbohydrate unit as ECA_PG_ attached to DAG through a pyrophosphate diester bond (75). This molecule is essential and has been implicated in insertion of proteins into the IM in a Sec-independent manner and in Sec-dependent protein translocation (74–80). Unlike ECA_PG_, the MPIase sugar subunit is built on DAG-PP with the first step being attachment of GlcNAc-P to DAG-P, a reaction catalyzed by CdsA and YnbB (76). The lipid donor for this reaction is CDP-DAG. The synthesis of MPIase has previously been shown to be independent of ECA synthesis (81) and our results confirm that these molecules use distinct lipid donors. The ECA genes necessary for biosynthesis of the ECA sugars and the formation of the ECA chain has also been shown to be dispensable for MPIase formation (81), although later results suggest they may contribute to some extent (82). Exploring the differences and similarities between these biosynthetic pathways will provide interesting insights into both pathways.

Our identification of PG as the lipid donor for ECA_PG_ biosynthesis provides an interesting parallel with lipoprotein synthesis: Lgt uses PG as the donor to transfer DAG to nascent lipoproteins (69). Despite this, PE is the most abundant phospholipid in the *E. coli* membrane. It is possible that these pathways use PG as the lipid donor due to the asymmetry of the IM, which contains more

PE in the inner leaflet than the outer leaflet (40). It is also possible that it is advantageous to the cell to use a less abundant membrane constituent for lipid modification as it can be more easily regulated, such as by decreasing PG levels in favor of CL during stationary phase (39, 73). Finally, the use of PG as a donor may be due to relative ease of recycling the glycerol head group compared to recycling of ethanolamine. These are intriguing questions for future study.

## MATERIALS AND METHODS

### Strains and growth conditions

The strains used in these experiments are listed in **Table 1**. Cultures were grown in LB Lennox at 37 °C unless otherwise noted. Where necessary for plasmid maintenance, media was supplemented with 20 mg/L chloramphenicol and/or 25 mg/L kanamycin. Where indicated, cultures were supplemented with 0.2% glucose, 0.2% arabinose, or 50 mM MgSO_4_. Unless otherwise noted, deletion alleles are from the Keio collection and were moved into our strains using *P1vir* transduction (83, 84). A deletion allele for *pgsA* was constructed using λ-Red recombineering and the primers listed in **Table 2** (85). Kanamycin resistance cassettes were flipped out using the Flp recombinase-FRT system as has been described (85). Professors Mikhail Bogdanov and William Dowhan (McGovern Medical School, University of Texas Houston) provided us with the kind gift of a previously published *pssA* disruption allele, *pss93::kan* (72). Strains with the *pss93::kan* allele were constructed by transforming strains with pDD72GM (86), which carries a wild-type *pssA* allele, transducing in the *pss93::kan* allele, and then curing the temperature sensitive pDD72GM plasmid.

**Table 1:** Strains used in this study.

| Strain | Genotyping | Reference | Plasmids and Alleles |
| --- | --- | --- | --- |
| MG1655 | K-12 F<sup>-</sup> $\lambda^-$ *rph-1* | (87) | |
| AM334 | MG1655 Δ*wecA* | (26) | |
| AM395 | MG1655 Δ*wzzE* Δ*waaL* | (26) | |
| AM1358 | MG1655 Δ*wzzE* Δ*waaL* Δ*clsA* Δ*clsB* Δ*clsC* | This study | |
| KM032 | MG1655 Δ*wzzE* Δ*waaL* *pss93::kan<sup>R</sup>* | This study | *pssA93::kan<sup>R</sup>* allele from (72) |
| AM1138 | MG1655 pCA24N | (56) | Plasmid is ASKA collection empty vector (88), cm<sup>R</sup> |
| AM1144 | MG1655 Δ*wzzE* Δ*waaL* pCA24N | This study | Plasmid is ASKA collection empty vector (88), cm<sup>R</sup> |
| AM1220 | MG1655 pCA24N-*cdsA* | This study | Plasmid from (88) |
| AM1286 | MG1655 Δ*wzzE* Δ*waaL* pCA24N-*cdsA* | This study | |
| AM1326 | MG1655 pCA24N-*pssA* | This study | Plasmid from (88) |
| AM1327 | MG1655 Δ*wzzE* Δ*waaL* pCA24N-*pssA* | This study | |
| AM1176 | MG1655 pCA24N-*psd* | This study | Plasmid from (88) |
| AM1283 | MG1655 Δ*wzzE* Δ*waaL* pCA24N-*psd* | This study | |
| AM1324 | MG1655 pCA24N-*pgsA* | This study | Plasmid from (88) |
| AM1325 | MG1655 Δ*wzzE* Δ*waaL* pCA24N-*pgsA* | This study | |
| AM1174 | MG1655 pCA24N-*pgpA* | This study | Plasmid from (88) |
| AM1282 | MG1655 Δ*wzzE* Δ*waaL* pCA24N-*pgpA* | This study | |
| AM1328 | MG1655 pCA24N-*pgpC* | This study | Plasmid from (88) |
| AM1329 | MG1655 Δ*wzzE* Δ*waaL* pCA24N-*pgpC* | This study | |
| AM1330 | MG1655 pCA24N-*clsA* | This study | Plasmid from (88) |
| AM1331 | MG1655 Δ*wzzE* Δ*waaL* pCA24N-*clsA* | This study | |
| AM1177 | MG1655 pCA24N-*clsB* | This study | Plasmid from (88) |
| AM1284 | MG1655 Δ*wzzE* Δ*waaL* pCA24N-*clsB* | This study | |
| AM1332 | MG1655 pCA24N-*clsC* | This study | Plasmid from (88) |
| AM1333 | MG1655 Δ*wzzE* Δ*waaL* pCA24N-*clsC* | This study | |
| KM014 | MG1655 Δ*wzzE* Δ*waaL* pCA24N pJW15 | This study | pJW15 is an empty *lux* reporter from (89); kan<sup>R</sup> |
| KM015 | MG1655 Δ*wzzE* Δ*waaL* pCA24N pJW15-P<sub>wec</sub> | This study | pJW15-P<sub>wec</sub> reporter plasmid (88) |
| KM011 | MG1655 $\Delta wzzE \Delta waaL$ pCA24N-*cdsA* pJW15-P<sub>wec</sub> | This study | |
| KM008 | MG1655 $\Delta wzzE \Delta waaL$ pCA24N-*pssA* pJW15-P<sub>wec</sub> | This study | |
| KM009 | MG1655 $\Delta wzzE \Delta waaL$ pCA24N-*psd* pJW15-P<sub>wec</sub> | This study | |
| KM005 | MG1655 $\Delta wzzE \Delta waaL$ pCA24N-*pgsA* pJW15-P<sub>wec</sub> | This study | |
| KM006 | MG1655 $\Delta wzzE \Delta waaL$ pCA24N-*pgpA* pJW15-P<sub>wec</sub> | This study | |
| KM012 | MG1655 $\Delta wzzE \Delta waaL$ pCA24N-*pgpC* pJW15-P<sub>wec</sub> | This study | |
| KM007 | MG1655 $\Delta wzzE \Delta waaL$ pCA24N-*clsA* pJW15-P<sub>wec</sub> | This study | |
| KM013 | MG1655 $\Delta wzzE \Delta waaL$ pCA24N-*clsB* pJW15-P<sub>wec</sub> | This study | |
| KM010 | MG1655 $\Delta wzzE \Delta waaL$ pCA24N-*clsC* pJW15-P<sub>wec</sub> | This study | |
| AM1162 | MG1655 pBAD33 | (56) | Plasmid from (65),<br>cm<sup>R</sup> |
| AM1158 | MG1655 $\Delta wzzE \Delta waaL$ pBAD33 | (56) | |
| AM1425 | MG1655 $\Delta pgpB \Delta pgpA$ pBAD33-*pgpC* $\Delta pgpC$ | This study | |
| AM1426 | MG1655 $\Delta wzzE \Delta waaL \Delta pgpB \Delta pgpA$ pBAD33-*pgpC* $\Delta pgpC$ | This study | |
| AM1407 | MG1655 pBAD33-*pgsA* $\Delta pgsA$ | This study | |
| AM1408 | MG1655 $\Delta wzzE \Delta waaL$ pBAD33-*pgsA* $\Delta pgsA$ | This study | |
| AM1409 | MG1655 $\Delta rcsF \Delta lpp$ pBAD33-*pgsA* $\Delta pgsA$ | This study | |
| AM1412 | MG1655 $\Delta rcsF \Delta lpp \Delta wzzE \Delta waaL$ pBAD33-*pgsA* $\Delta pgsA$ | This study | |
| AM1413 | MG1655 $\Delta clsA \Delta clsB$ pBAD33-*clsA* $\Delta clsC$ | This study | |
| AM1414 | MG1655 $\Delta wzzE \Delta waaL \Delta clsA \Delta clsB$ pBAD33-*clsA* $\Delta clsC$ | This study | |

**Table 2:**
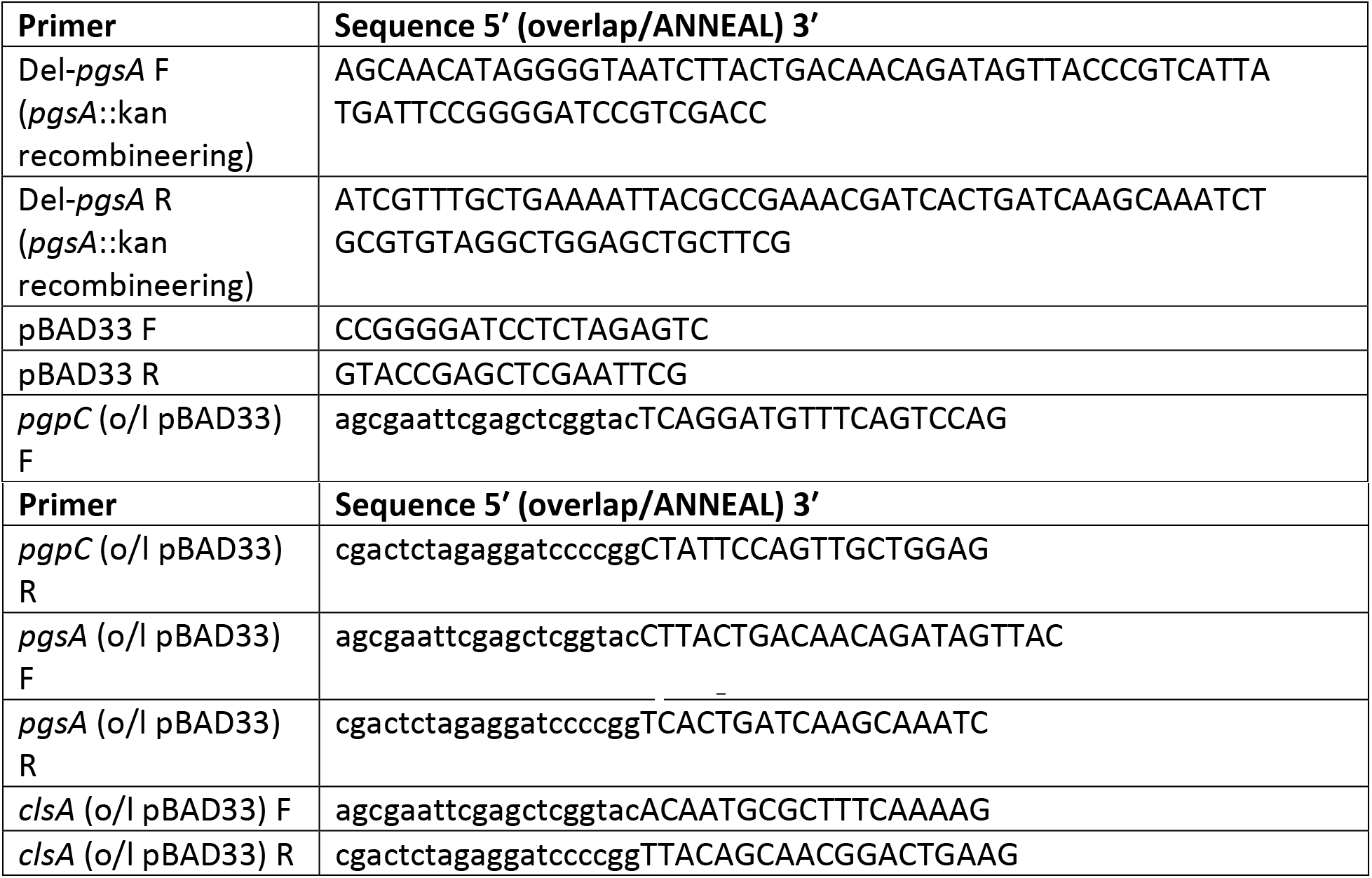
Primers used in this study.

IPTG inducible overexpression constructs were derived from the ASKA collection of *E. coli* gene overexpression plasmids and were transformed into the indicated strains using standard molecular biology techniques. For construction of PgsA, PgpC, and ClsA depletion strains, *pgsA*, *pgpC*, and *clsA* were cloned into P_BAD_33 using Gibson Assembly (65). Briefly, P_BAD_33 was linearized by PCR using the P_BAD_33 F and R primers (**Table 2**). The inserts amplified from MG1655 genomic DNA using the primers listed in **Table 2** from 30 bp upstream of the ORF to the end of the ORF. The PCR fragments were assembled using HiFi Assembly Master Mix (New England Biolabs) as per the manufacturer’s instructions.

### Quantification of ECA levels

ECA_LPS_ levels were quantified by staining cells with Alexafluor 488-conjugated WGA (Thermo Fisher Scientific) as we have described (56). Briefly, 250 μL aliquots of washed overnight cultures were combined with 10 μg/mL WGA in PBS and were incubated 10 minutes at room temperature. Cells were washed twice and resuspended in PBS. Fluorescence (Excitation 485 nm, Emission 519 nm) and OD_600_ were read in a BioTek Synergy H1 plate reader. Fluorescence relative to OD was calculated.

ECA_PG_ was detected through immunoblot analysis as we have described (26, 56). Briefly, samples from overnight cultures were normalized to equal OD_600_ and resuspended in BugBuster Protein Extraction Reagent (Millipore Sigma) followed by an equal volume of Laemmli Sample Buffer (BioRad). Samples were run on 12% acrylamide gels and transferred to nitrocellulose. Blots were probed with α-ECA antibody at a 1:30,000 dilution and a goat α-rabbit-HRP secondary antibody (Prometheus) at a 1:100,000 dilution. α-ECA antibody was a kind gift of Professor Renato Morona (University of Adelaide). Detection was performed using Prosignal Pico ECL (Prometheus) using Prosignal ECL-blotting film (Prometheus).

### Luciferase reporter assays

Activity of the P*_wec_* promoter was assayed with a plasmid-based *P_wec_:luxCDABE* reporter as we have described (56) with minor modifications. Cultures were grown overnight in media without IPTG. For reporter assays, strains were sub-cultured 1:100 into media with the indicated concentration of IPTG. Luminescence and the OD_600_ were assayed in technical quadruplicate in a BioTek Synergy H1 plate reader every 3 minutes for 5 hours. Luminescence relative to OD_600_ was calculated and technical replicates were averaged. Then, the fold value of each sample to the empty vector control for each time point was calculated.

### Depletion growth curves

For depletion experiments, the indicated strains were grown overnight with arabinose. Then, cultures were washed once in plain LB and diluted 1:500 or 1:10,000, as indicated, into media with either arabinose to induce expression from the P_BAD_ plasmid or glucose to repress expression. Cultures were transferred to a 24-well plate and were incubated shaking at 37 °C in a BioTek Synergy H1 plate reader. OD_600_ was assayed every 5 minutes for 12 hours.

## ACKNOWLEDGEMENTS

We thank members of the Mitchell lab for productive discussions and for critical reading of our manuscript. We thank Professor Tracy Raivio (University of Alberta) for the pJW15 plasmid and Professor Renato Morona (University of Adelaide) for the ECA antibody. We thank Professors Mikhail Bogdanov and William Dowhan (McGovern Medical School, University of Texas Houston) for providing us with strains carrying a *pssA* disruption and complementing plasmid. The work described here was supported by the National Institute of Allergy and Infectious Disease under award number R01-AI155915 (to A.M.M.).

